# Sensory neuron-associated macrophages proliferate in the sensory ganglia after peripheral nerve injury in a CX3CR1 signaling dependent manner

**DOI:** 10.1101/2022.03.22.485276

**Authors:** Rafaela M. Guimarães, Conceição E. A. da Silva, Marcela Davoli-Ferreira, Francisco Isaac F. Gomes, Atlante Mendes, Miriam M. Fonseca, Samara Damasceno, Larissa P. Andrade, Fernando Q. Cunha, José C. Alves-Filho, Thiago M. Cunha

## Abstract

Resident macrophages are distributed across all tissues and are highly heterogeneous as a consequence of adaptation to different tissue-specific environments. The resident macrophages of the sensory ganglia (sensory neuron-associated macrophages, sNAMs) are in close contact with the cell body of primary sensory neurons and might play physiological and pathophysiological roles. After peripheral nerve injury, there is an increase in the population of macrophages in the sensory ganglia which have been involved in different conditions, especially in neuropathic pain development and nerve regeneration. However, it is still under debate whether macrophages accumulation in the sensor ganglia after peripheral nerve injury is due to the local proliferation of resident macrophages or as a result of blood monocytes infiltration. Here, we confirmed that the number of macrophages increased in the DRGs after spared nerve injury (SNI) model in mice. By using different approaches, we found that the increase in the number of macrophages in the DRGs after SNI is mainly in consequence of the proliferation of resident CX3CR1^+^ macrophages but not due to infiltration of CCR2^+^ blood monocytes. These proliferating macrophages are the source of the production of pro-inflammatory cytokines such as TNF and IL-1b. In addition, we found that CX3CR1 signaling is involved in the sNAMs proliferation after peripheral nerve injury. In summary, these results indicated that peripheral nerve injury leads to sNAMs proliferation in the sensory ganglia in a CX3CR1-dependent manner. In conclusion, sNAMs proliferation could be modulated to change pathophysiological conditions such as chronic neuropathic pain and/or nerve regeneration.

## 1. Introduction

Most organs across the body contain tissue-resident populations of macrophages (Wynn, Chawla, and Pollard 2013; Perdiguero and Geissmann 2016). Historically, these resident cells are well-known for participating in host defense as well as in the clearance of dead cells and tissue debris, whereby they contribute to a range of pathological processes (Ginhoux and Guilliams 2016; Davies et al. 2013; Wynn and Vannella 2016). It has been appreciated that beyond their classical role in tissue inflammation, macrophages are a heterogeneous population, exhibiting high functional plasticity that is correlated with the specific functions of each tissue and niche they reside (Epelman, Lavine, and Randolph 2014; Van Hove et al. 2019).

Among the distinct subsets of tissue-resident macrophages, those that colonize the peripheral nervous system (PNS) have only recently been studied in more detail. In the PNS, macrophages distributed along the sciatic nerve and in the sensory ganglia (e.g. dorsal root ganglia - DRG and trigeminal ganglion) are in close contact with the primary sensory neurons, known as sensory neurons-associated macrophages (sNAMs) (C.E.A. Silva, Guimarães, and Cunha 2021; Kolter, Kierdorf, and Henneke 2020). These resident macrophages seem to play a crucial role in nervous tissue repair and in the development of neuropathic pain caused by peripheral nerve injury (Wang et al. 2020; Iwai et al. 2021).

In the injured peripheral nerves, the sNAMs functions are associated with phagocytosis of cell debris and release of early inflammatory mediators, which in turn contribute to the recruitment of neutrophils and CCR2-expressing monocytes (Calvo, Dawes, and Bennett 2012; C.F. Kim and Moalem-Taylor 2011; Lindborg, Mack, and Zigmond 2017; Jung et al. 2009). These changes propagate the neuroinflammatory response at the level of sensory ganglia characterized by activation of glial cells and sNAMs and the consequent release of pro-inflammatory cytokines, in particular, IL-6, IL-1β, and TNF (Yu et al. 2020; Kwon et al. 2013; Ji, Xu, and Gao 2014).

Several studies using different peripheral nerve injury models have described an increase in the number of macrophages/monocytes surrounding the cell body of sensory neurons in the DRGs (Kwon et al. 2013; Liu et al. 2010; D. Kim, You, et al. 2011; Huang et al. 2014; Kallenborn-Gerhardt et al. 2014; Luo et al. 2019). Although these studies have referred to this increase as a result of cell infiltration, it is still unclear whether it would be in view of local resident cells proliferation or infiltration of inflammatory monocytes.

Despite the complexity of determining macrophage populations, the high expression of classical markers such as CX3CR1 (fractalkine receptor) and CCR2 is often used to distinguish yolk sac-derived embryonic macrophages and bone marrow (BM)-derived monocytes, respectively (Stremmel et al. 2018; Geissmann, Jung, and Littman 2003). In fact, CX3CR1^+^ macrophages are considered a long-lived subset patrolling in the blood and mainly expressed in tissue-resident macrophages, while the CCR2^+^ monocytes are a short-lived subset that is actively recruited into the inflamed tissue (Yona et al. 2013; Auffray et al. 2007). In the present study, by using a combination of different approaches, we addressed the question of whether the origin of the injury-induced macrophages expansion in the DRG depends on the blood circulating monocytes infiltration or the resident macrophages proliferation. Briefly, our findings indicate that after peripheral nerve injury, the raise in the macrophages number in the sensory ganglia (e.g. DRGs) is essentially due to the proliferation of local/resident CX3CR1^+^ macrophages than the recruitment of CCR2^+^ blood circulation monocytes. Furthermore, we showed that CX3CR1 signaling on NAMs mediates their proliferation/activation triggering pro-inflammatory cytokines synthesis.

## 2. Results

### SNI triggers an increase in the number of macrophages in the DRGs

There is evidence showing that peripheral nerve injury induces an increase in the number of macrophages in the sensory ganglia (Yu et al. 2020; Kwon et al. 2013). Herein we sought to characterize this process using a classical model of peripheral nerve injury, spared nerve injury (SNI) in mice (Decosterd and Woolf 2000). Firstly, after nerve injury we evaluated the gene expression profile of macrophage markers, such as *Aif1* (IBA-1) and *Csf1r* (CSFR1), in the ipsilateral DRGs at different time points (Fig. 1A). We found that *Aif1* and *Csf1r* expression were up regulated from day 3, reaching a peak between 7 and 10 days after SNI induction (Fig. 1B). Additionally, immunofluorescence analyses also revealed an increased number of macrophages (IBA-1^+^ cells) in the DRGs after 7 days of nerve lesion (Fig. 1C-D). Using flow cytometry, we confirmed a rise in the macrophages CD11b^+^Ly6G^-^ cells number in the DRGs at 7 days after SNI induction (Fig. S1A). Together, these data confirmed that peripheral nerve injury macrophage population increases in the sensory ganglia.

**Figure 1.**
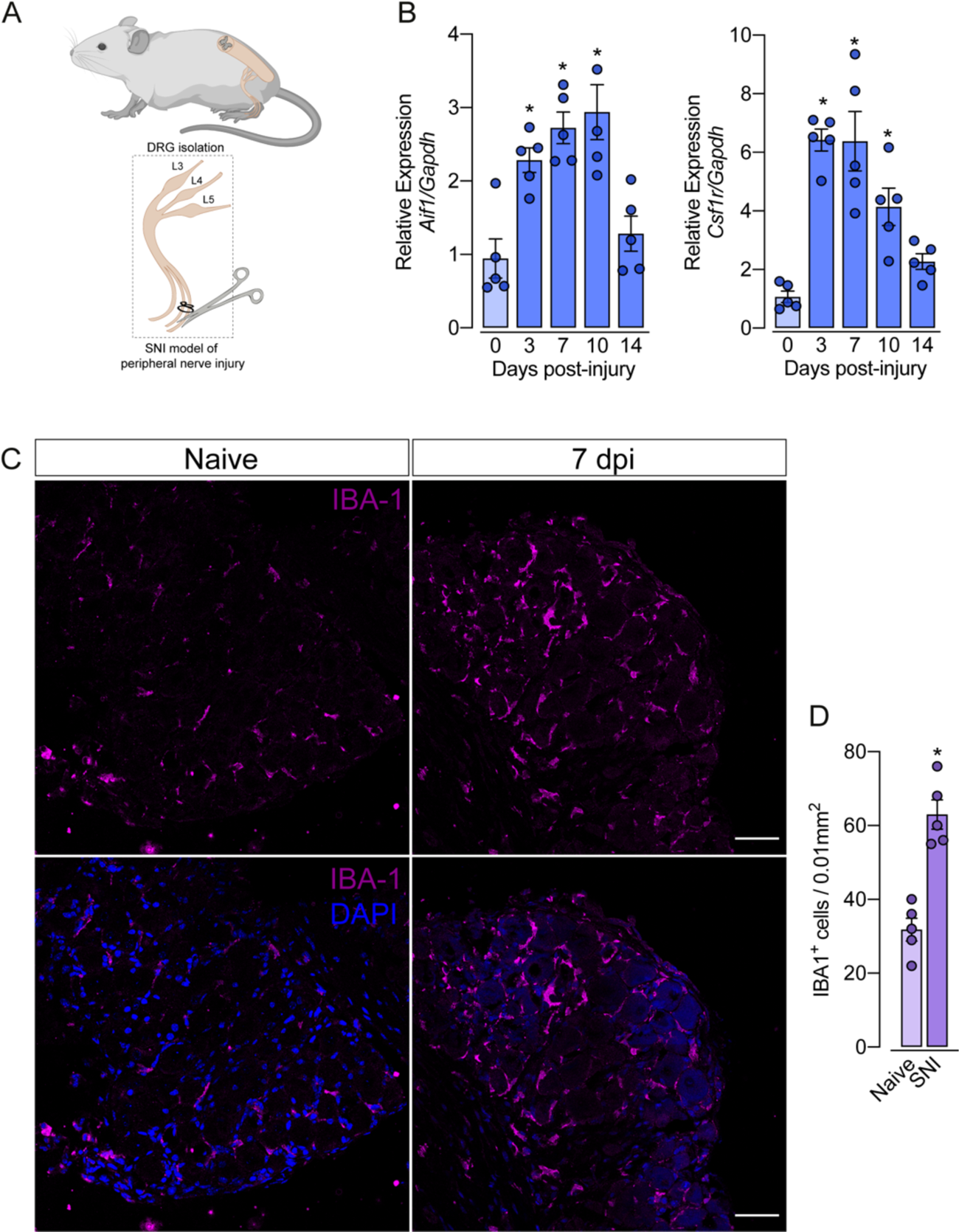
Spared nerve injury (SNI) model induces an increase in the number of macrophages in the DRG. **(A)** Schematic representation of the experimental design of SNI induction in wild type mice, showing the sciatic nerve, its branches and the dorsal root ganglia (DRG; L3, L4 and L5) harvested. **(B)** Time course of *Csf1r* and *Aif1* mRNA expression relative to *Gapdh* in the DRGs after SNI or naive mice (n = 4-5). **(C)** Representative confocal images of the DRG from WT mice at 7 days after SNI (n = 3). **(D)** Quantification of macrophages (IBA**^+^** cells) in L4 DRGs at 7 days after SNI (n = 5). Error bars show the mean ± S.E.M. P values were determined by one-way ANOVA followed by Bonferroni’s post hoc test. *, P < 0.05. Data are representative of at least 2 independent experiments.

### CX3CR1**^+^** macrophages population expand in the DRGs after peripheral nerve injury independently of CCR2**^+^** monocytes

As mentioned previously, it is still unclear whether circulating monocytes can infiltrate into the sensory ganglia (e.g. DRG) and might account for the increase in the number of macrophages observed after peripheral nerve injury. To address this question, we employed a series of different experimental approaches. Firstly, we used *Cx3cr1^GFP/+^/Ccr2^RFP/+^* reporter mice, which might be useful to distinguish the typical CX3CR1^+^ tissue-resident macrophages from CCR2^+^ blood monocytes (Fig. 2A). Notably, while the number of CX3CR1^+^ cells in the DRGs increased after SNI induction (Fig. 2B-C), CCR2^+^ inflammatory monocytes numbers did not change (Fig. 2B-C), indicating that blood CCR2^+^ monocytes did not infiltrate into the sensory ganglia after peripheral nerve injury. Corroborating these data, we found an increase in the expression of *Cx3cr1* in the DRG after SNI, and flow cytometry analysis confirmed a significant increase in the number of CX3CR1^+^ cells when compared to the naive group (Fig. S2A-B). As a positive control of CCR2^+^ monocytes infiltration into the DRGs, we used a murine model of HSV-1 infection (J.R. Silva et al. 2017). As expected, we observed a significant CCR2^+^ blood monocytes infiltration in the DRG of HSV-1-infected mice (Fig. S3A-B). Additionally, by using deficient mice for the chemokine receptor CCR2 (*Ccr2*^−/−^ mice), we noticed that the increase of *Cx3cr1 gene* expression after SNI did not change in these mice when compared to *Ccr2*-sufficient animals (Fig. S3C). These results suggest that infiltration of CCR2^+^ monocytes is not required for the expansion of CX3CR1^+^ resident-macrophages in the DRGs after peripheral nerve surgery.

**Figure 2.**
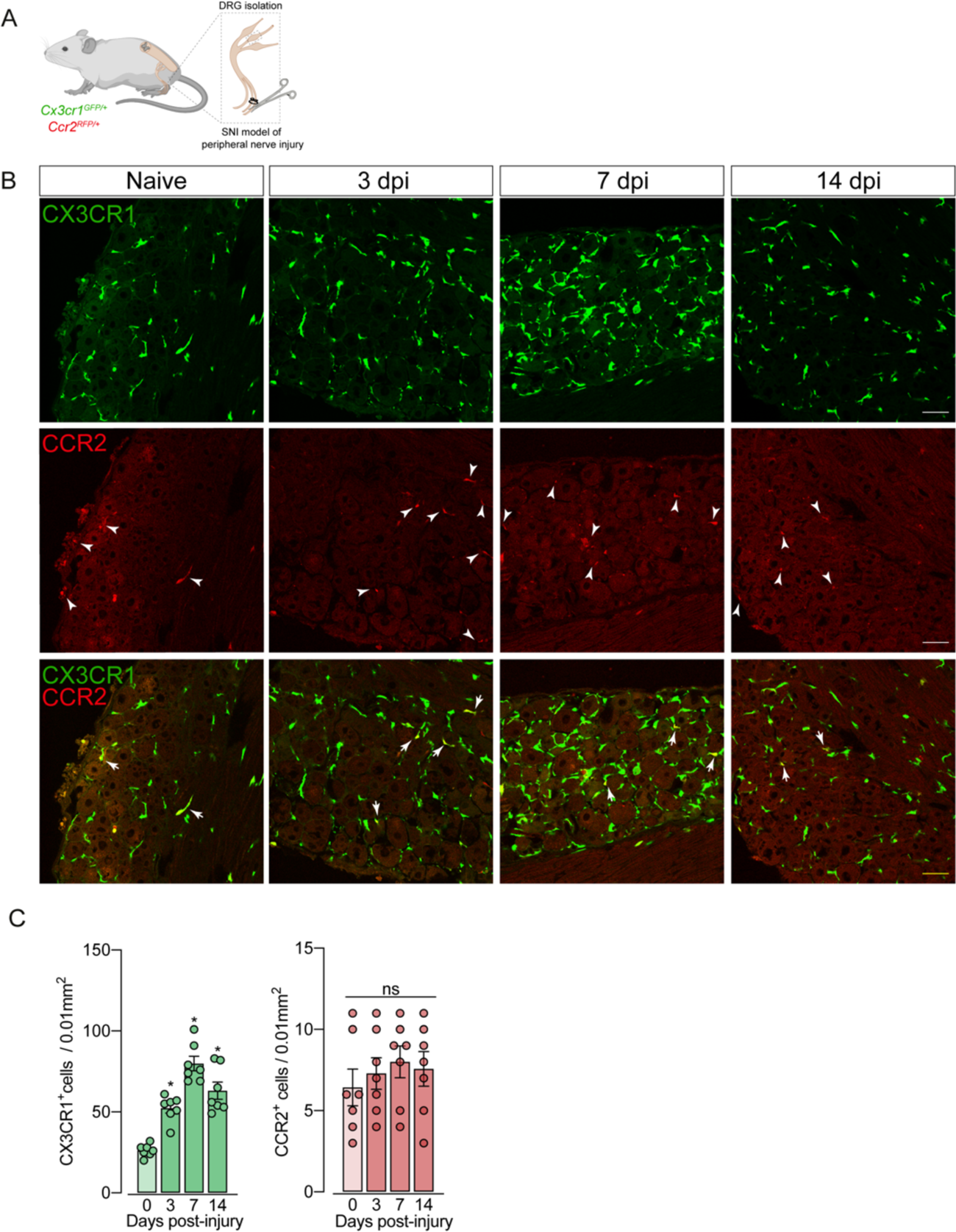
SNI induces an increase of CX3CR1^+^ macrophages, but not of CCR2^+^ monocytes in the DRGs. (**A**) Schematic representation of the experimental design of SNI induction in *Cx3cr1^GFP/+^/Ccr2^RFP/+^* mice. **(B)** Representative confocal images of the DRG from *Cx3cr1^GFP/+^/Ccr2^RFP/+^* mice 3, 7- and 14-days post-injury (dpi). CX3CR1-GFP**^+^** macrophages are shown in green and CCR2-RFP**^+^** monocytes are shown in red (indicated white arrows). Scale bars: 50 μm. **(C)** Quantification of macrophages (CX3CR1-GFP**^+^** cells) and monocytes (CCR2-RFP**^+^** cells) in L4 DRGs at 0, 3, 7 and 14 dpi (n = 7). Error bars show the mean ± S.E.M. P values were determined by one-way ANOVA followed by Bonferroni’s post hoc test. *, P < 0.05; ns, not significant. Data are representative of at least 3 independent experiments.

To support these data, next, we performed parabiosis as an additional approach to elucidate the possible infiltration of blood-borne monocytes in the DRG after SNI. Parabiotic WT and GFP pairings mice were surgically joined and remained sharing circulation for 4 weeks, followed by the SNI induction (Fig. 3A-B). In agreement with our earlier results, the number of macrophages (IBA-1^+^ cells) increased in the DRG of WT mice 7 days after SNI. However, the number of GFP^+^ cells in the DRGs of WT mice remained constant in both naïve and SNI groups (Fig. 3C-D). Next, using the same experimental approach, we performed parabiosis in WT and *Cx3cr1^GFP/+^/Ccr2^RFP/+^* mice (Fig. 4A-B). Despite the increase of macrophages (IBA-1^+^ cells) in the DRG after SNI induction, we could not detect CCR2^+^ or CX3CR1^+^ cells (Fig. 4C-D). Altogether, these findings strongly suggest that the increase in the number of macrophages in the DRGs after peripheral nerve injury is likely to result from the expansion/proliferation of CX3CR1^+^ resident macrophages regardless of infiltration of peripheral blood CCR2^+^ monocytes.

**Figure 3.**
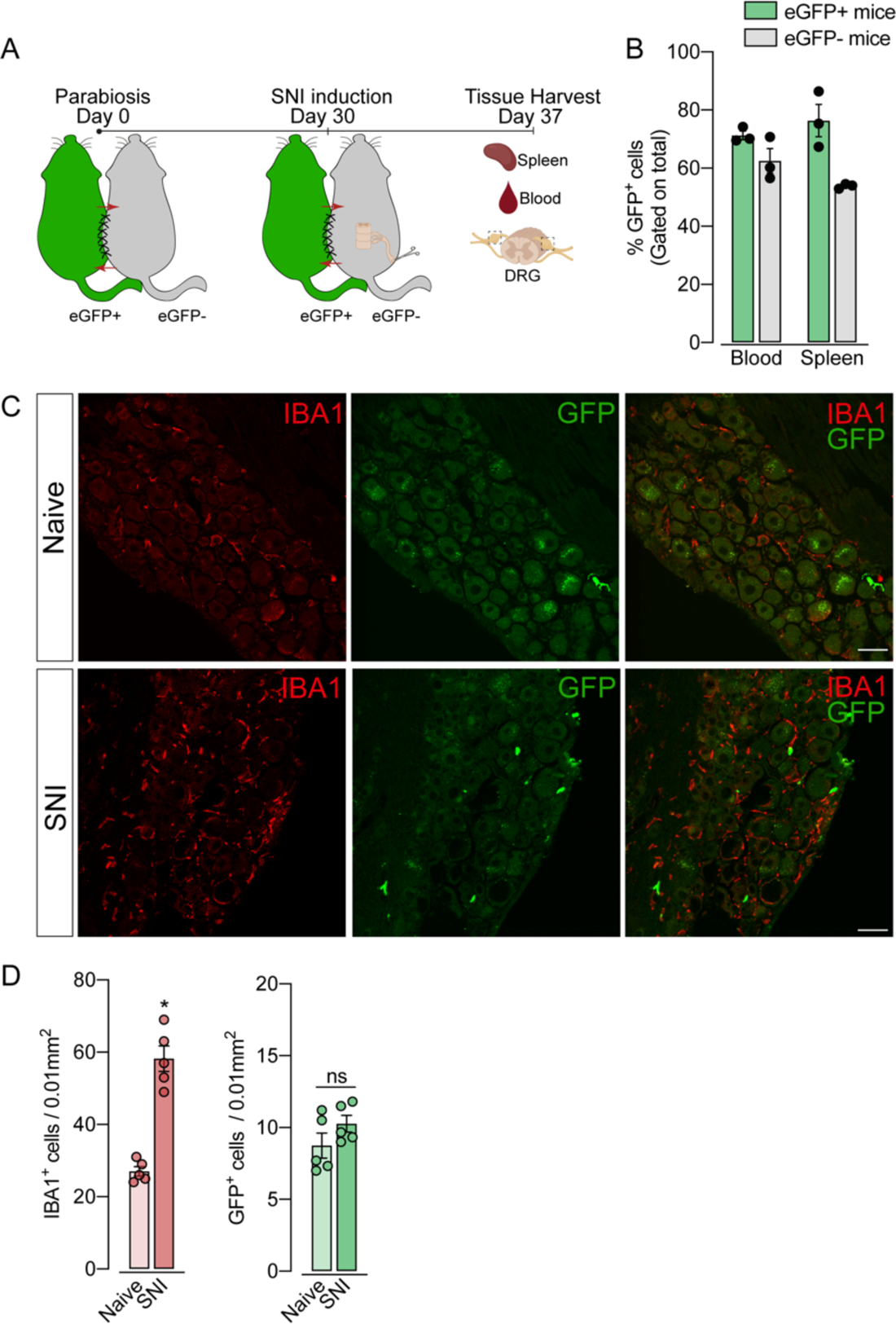
Parabiosis reveals no infiltration of blood leukocytes in the DRGs after SNI. **(A)** Schematic representation of parabiotic mouse pairs (eGFP**^+^** and eGFP-mice). After 30 days, the SNI model was induced in eGFP-mice and maintained for 7 days, and then, spleen, blood and DRGs were harvested. **(B)** Frequency of GFP**^+^** cells in the blood and spleen from eGFP**^+^** and eGFP**^-^** mice after 37 days of parabiosis. **(C)** Representative confocal images of the DRG from naïve or SNI eGFP-mice. GFP**^+^** cells are shown in green and IBA1**^+^** cells are shown in red. Scale bars: 50 μm. **(D)** Quantification of GFP**^+^** and IBA1**^+^** cells staining in naïve or ipsilateral L4 DRGs (n = 5 pairs of mice). Data are representative of two independent experiments. Error bars show mean ± SEM. P values were determined by two-tailed Student’s *t* test. *, P < 0.05; ns, not significant. Data are representative of at least 2 independent experiments.

**Figure 4.**
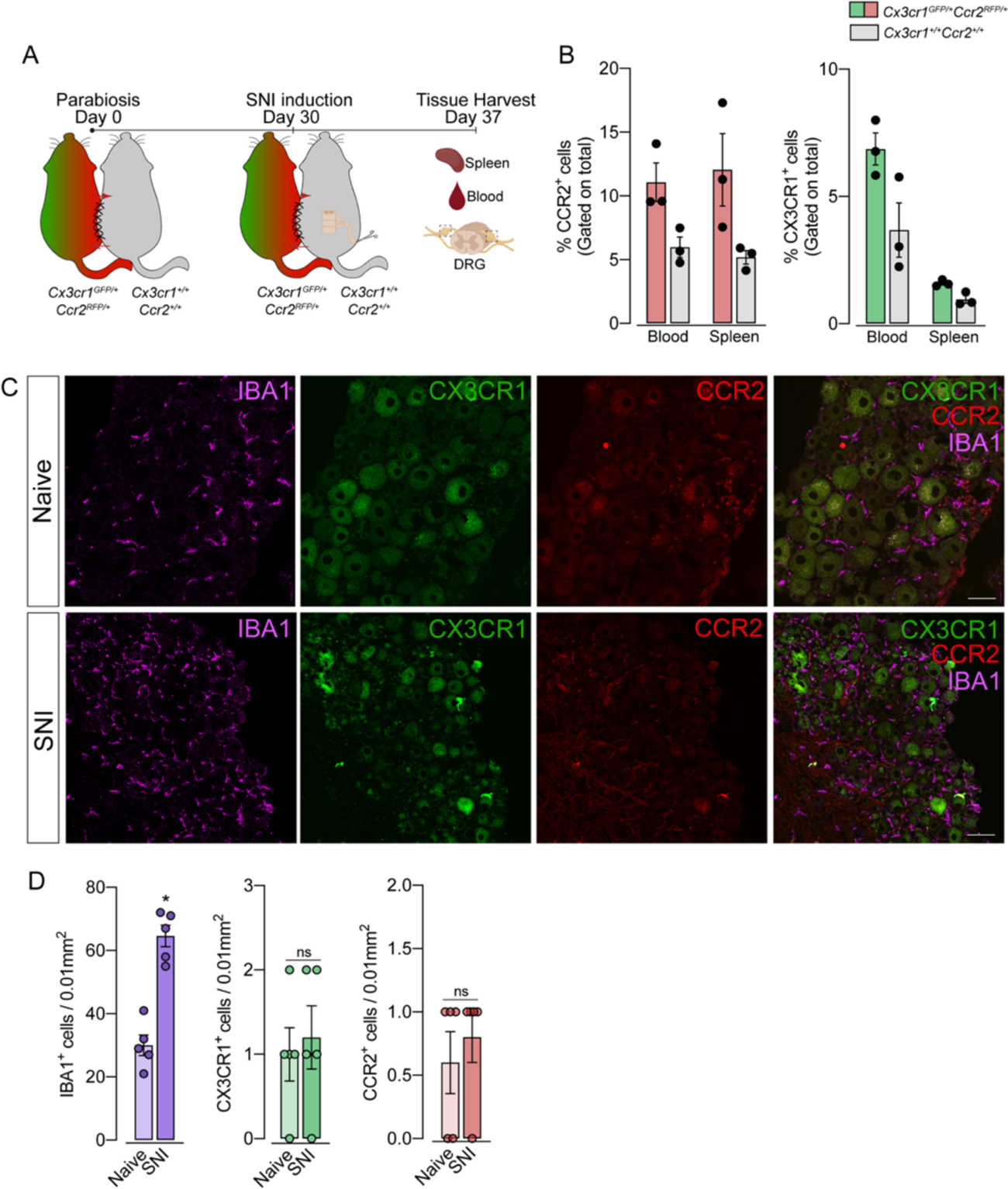
Parabiosis reveals no infiltration of blood CX3CR1^+^ or CCR2^+^ cells in the DRG after SNI. **(A)** Schematic representation of parabiotic mouse pairs *Cx3cr1^GFP/+^/Ccr2^RFP/+^* and *Cx3cr1^+/+^/Ccr2^+/+^* mice joined for 37 days. After 30 days, the SNI model was induced in *Cx3cr1^+/+^/Ccr2^+/+^* mice and maintained for 7 days, and then, spleen, blood and DRG were harvested. **(B)** Frequency of CX3CR1**^+^** cells and CCR2**^+^** cells in the blood and spleen from parabiotic mice. **(C)** Representative confocal images of the DRG from naïve or SNI *Cx3cr1^+/+^/Ccr2^+/+^* mice. CX3CR1**^+^** cells are shown in green, CCR2**^+^** cells are shown in red and IBA1**^+^** cells are shown in magenta. Scale bars: 50 μm. **(D)** Quantification of IBA1**^+^** cells staining in naïve or ipsilateral L4 DRGs (n = 5 pairs of mice). Data are representative of two independent experiments. Error bars show mean ± SEM. P values were determined by two-tailed Student’s *t* test. *, P < 0.05; ns, not significant. Data are representative of at least 2 independent experiments.

### CX3CR1**^+^** resident macrophages proliferated in the sensory ganglia after peripheral nerve injury

Given the increase in the number of macrophages in the DRGs after SNI in the absence of circulating monocytes infiltration, we hypothesized that CX3CR1^+^ resident macrophages in the DRG could undergo rapid and local proliferation after peripheral nerve injury. Corroborating this hypothesis, we found that the expansion of CX3CR1^+^ macrophages in the DRGs was accompanied by an increase in Ki67^+^ staining in CX3CR1^+^ macrophages three days after SNI indicating a high proliferation profile (Fig. 5A-B). These results indicate that the CX3CR1^+^ resident macrophage population expands in the DRGs and is also likely to account for the increase in the number of macrophages in the sensory ganglia observed after peripheral nerve injury.

**Figure 5.**
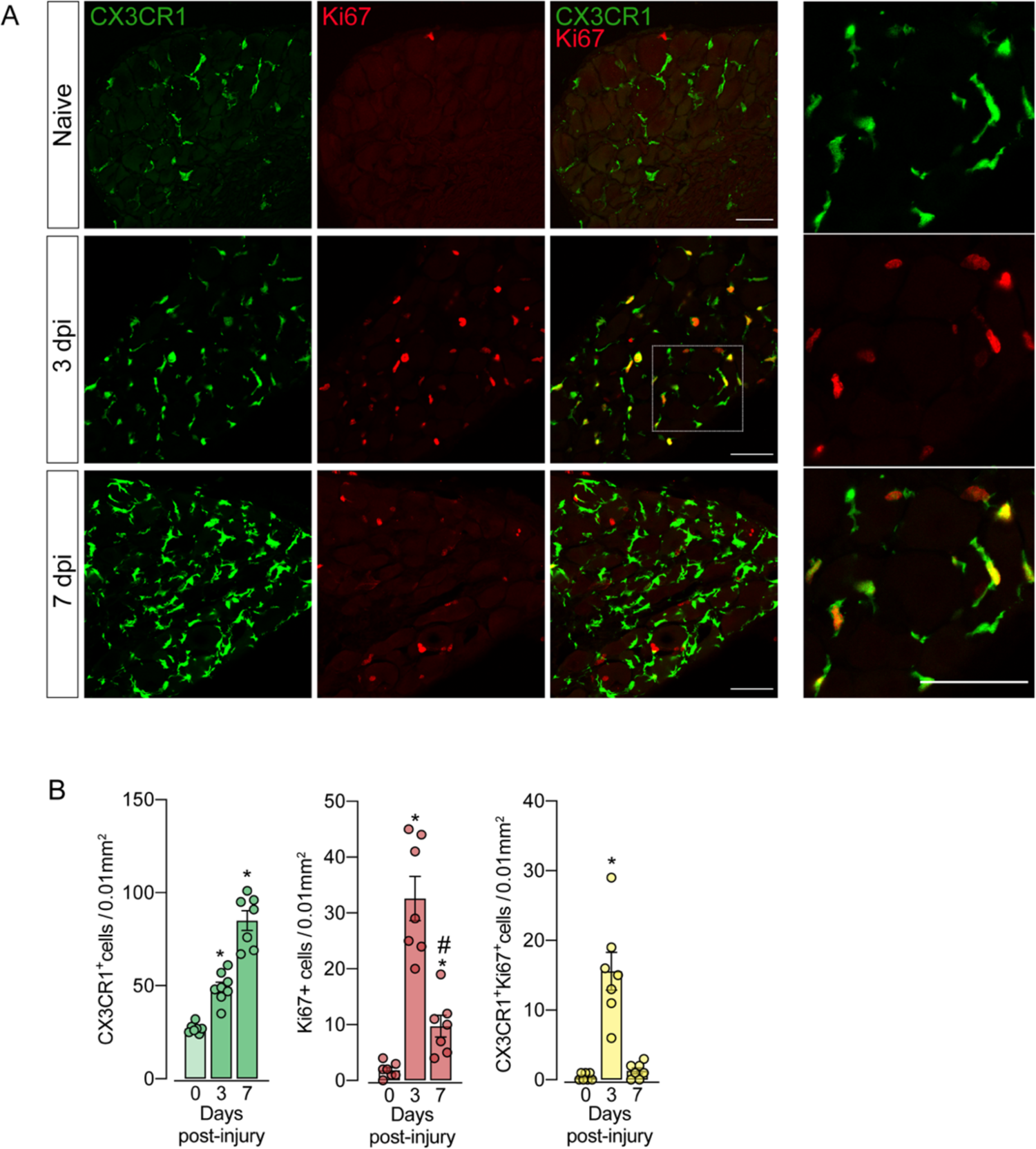
sNAMs proliferate in the sensory ganglia after peripheral nerve injury. **(A)** Representative confocal images of the DRG from *Cx3cr1^GFP/+^* mice after 3 and 7 days of SNI induction. CX3CR1**^+^** cells are shown in green and Ki67**^+^** cells are shown in red. Dotted boxes show regions of higher magnification in the DRG. Scale bars: 50 μm. (**B**) Quantification of CX3CR1^+^, Ki67^+^ and CX3CR1^+^Ki67^+^ cells (% of CX3CR1-proliferating cells) in naïve or ipsilateral L4 DRGs (n = 7-8). Error bars show the mean ± SEM. P values were determined by one-way ANOVA followed by Bonferroni’s post hoc test. *, P < 0.05. Data are representative of at least 3 independent experiments.

### CX3CR1 signaling mediates sNAMs expansion in the DRG after peripheral nerve injury

After characterizing the proliferation of resident macrophages in the sensory ganglia triggered by peripheral nerve injury, we sought to investigate possible mechanisms involved. Since CX3CR1 is expressed in sNAMs (Wang et al. 2020; Kolter et al. 2019; Chakarov et al. 2019) and there is evidence that CX3CR1 signaling mediates microglia proliferation in the spinal cord after peripheral nerve injury (Gu et al. 2016; Suter et al. 2009; Rotterman and Alvarez 2020), we tested whether this signaling would be involved on SNI-induced sNAMs expansion. For this purpose, we took advantage of *Cx3cr1^GFP/GFP^* mice (*Cx3cr1 null mice*) and evaluated the increase in the number of macrophages in the DRGs after SNI compared to *Cx3cr1^GFP/+^* mice (CX3CR1-sufficient mice). By using *Cx3cr1^GFP/GFP^*, we explore whether the CX3CR1 receptor contributed to the expansion of DRG resident macrophages. Interestingly, *Cx3cr1^GFP/GFP^* mice, at steady-state, did not show any difference in the frequency and number of CX3CR1^+^CD11b^+^ cells in the sensory ganglia (DRGs L3-L5) compared to heterozygous littermate controls (*Cx3cr1^GFP/+^* mice). Nevertheless, the increase of CX3CR1^+^CD11b^+^ cell numbers observed after SNI was reduced in the absence of the CX3CR1 (Fig. 6A-B). These results suggest that CX3CR1 signaling does not control the survival/seeding of sNAMs in the sensory ganglia, but might be important for their proliferation after peripheral nerve injury.

**Figure 6.**
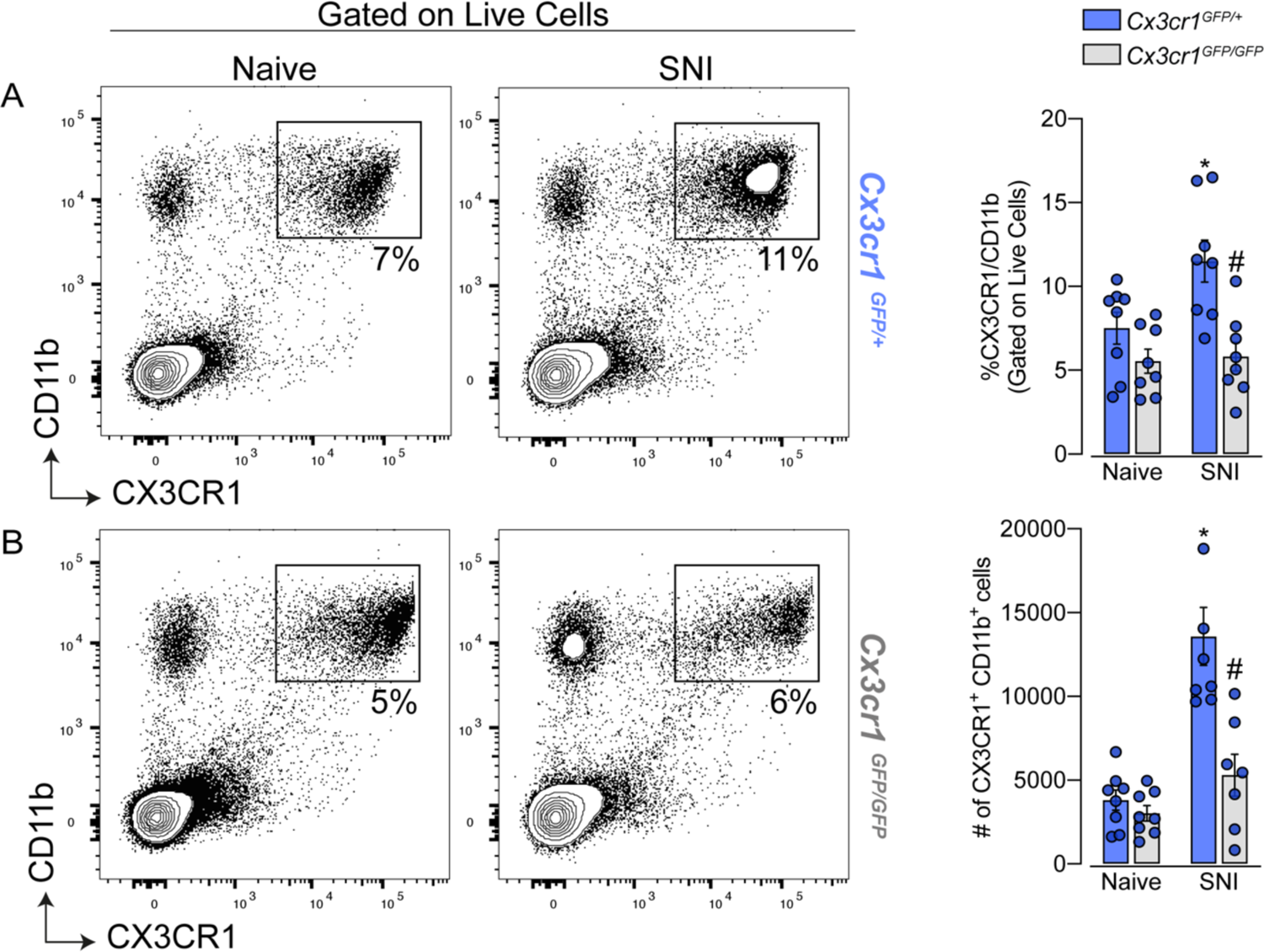
Cx3cr1 signaling involves macrophage expansion in the DRG after peripheral nerve injury. **(A-B)** Frequency and absolute number of CD45^+^ and CX3CR1^+^CD11b^+^ cells in the DRG from *Cx3cr1^GFP/+^* or *Cx3cr1^GFP/GFP^* mice at 7 days after SNI analyzed by flow cytometry (n = 7-8). Error bars show the mean ± S.E.M. P values were determined by one-way ANOVA followed by Bonferroni’s post hoc test. *, P < 0.05. Data are representative of at least 3 independent experiments.

### sNAMs are the main source of pro-inflammatory cytokines production after peripheral nerve injury: role of CX3CR1 signaling

After peripheral nerve injury, the expression of cytokines increased in the sensory ganglia, which are involved in the pathophysiological processes such as neuropathic pain. Among these cytokines, TNF, IL-1b, and IL-6 seem to be the most important ones (Yu et al. 2020; Kwon et al. 2013; Ji, Xu, and Gao 2014). Nevertheless, the cellular source of these cytokines in the sensory ganglia after peripheral nerve injury is still controversial. Herein, we confirmed this evidence and observed an up-regulation of *Tnf, Il1b,* and *Il6* mRNA in the DRGs of mice 7 days after SNI compared to naive-control mice (Fig. 7A). In an attempt to identify the cellular source of these cytokines, initially, we took advantage of publicly available single-cell RNAseq data from DRGs cells (Avraham et al. 2020). After the re-analyze of this data, we were able to identify 12 different cellular clusters, including sNAMs in the DRGs (Fig 7B). In addition, the expression of *Tnf and Il1b* after peripheral nerve injury was confined in the sNAMs cluster (Fig. 7B), whereas the expression of *Il6* was not conclusively defined (Fig. 7B). In order to confirm these data, next we performed cell sorting of both CD45**^-^** and CX3CR1^+^CD11b^+^ cells from DRGs from *Cx3cr1^GFP/+^* mice ipsilateral and contralateral to the SNI injury. We found that *Tnf and Il1b* gene expression is only detected in CX3CR1^+^CD11b^+^ cells harvested from naïve animals and their expression only increased in this specific cell population after SNI induction (Fig. 7C). On the other hand, *Il6* expression was detected mainly on CD45^-^ cells (Fig. 7C). Finally, we sought to investigate whether the increase in the production of pro-inflammatory cytokines after peripheral nerve injury is also dependent on CX3CR1 signaling. Whereas the increase of *Tnf and Il1b* in the DRGs from WT mice after SNI, it was reduced in DRGs from *Cx3cr1^GFP/GFP^* mice (Fig. 7D). On the other hand, *Il6* expression increased in the DRGs similarly in both mice genotypes (Fig. 7D). Altogether, these results indicated that sNAMs are the main source of CX3CR1 signaling-dependent pro-inflammatory cytokines (e.g. IL-b and TNF) in the sensory ganglia after peripheral nerve injury, whereas IL-6 seems to be induced mainly in non-immune cells.

**Figure 7.**
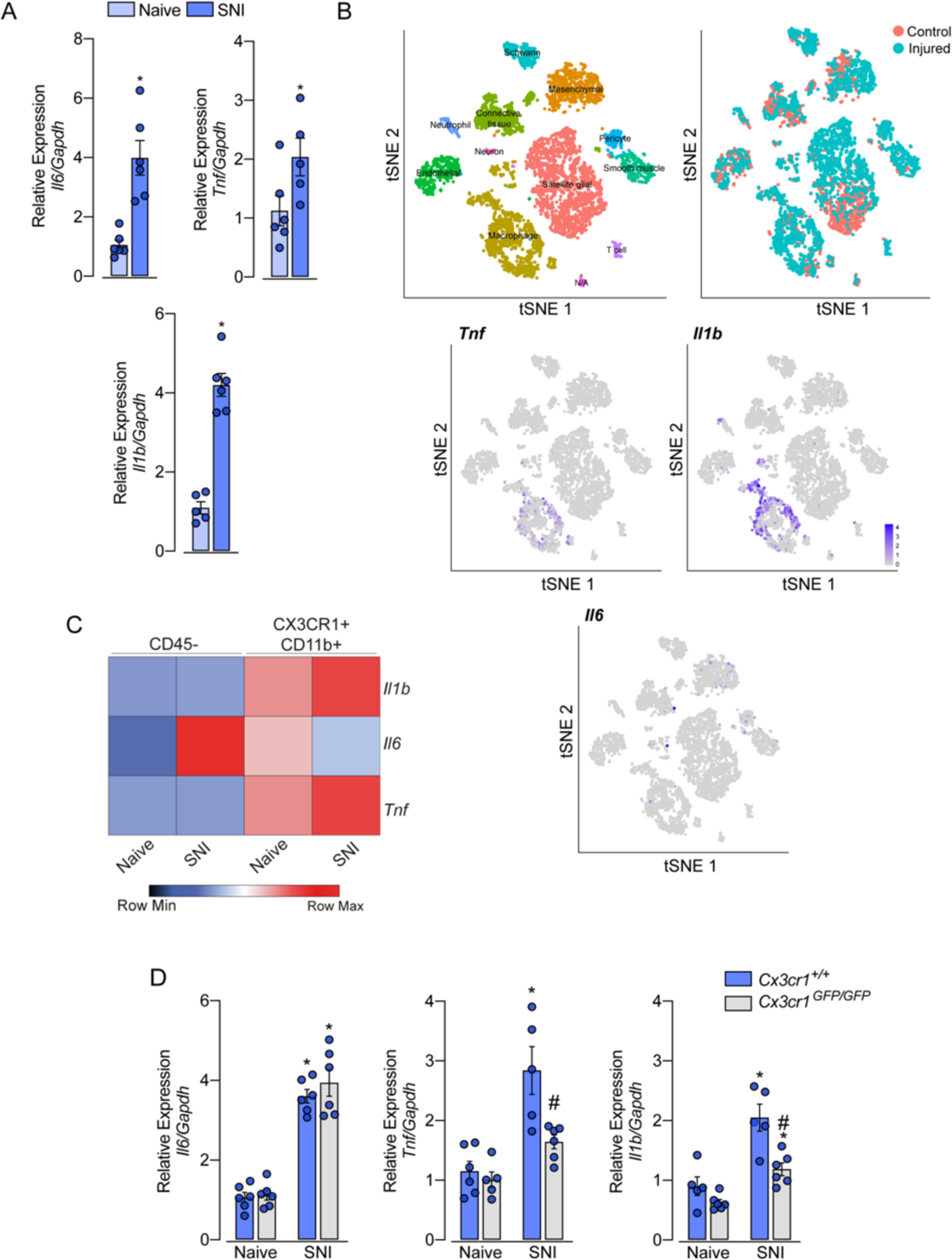
Role of sNAMs in the production of pro-inflammatory cytokines in the sensory ganglia after SNI. (**A**) qPCR analysis of *Il6, Il1b* and *Tnfa* mRNA expression relative to *Gapdh* levels in the DRG at 7 days after SNI or naive mice (n = 5-6). **(B)** t-SNE plot analysis showing clusters of cell populations and expression profile of *Tnfa, Il1b and Il6* in the DRGs from naive and after peripheral nerve injured mice. **(C)** qPCR analysis of *Il6*, *Il1b* and *Tnfa* mRNA expression relative to *Gapdh* levels in CD45^-^ or CX3CR1^+^CD11b^+^ cells isolated from DRGs at 7 days after SNI or naïve *Cx3cr1^GFP/+^* mice (n = 6 pooled). **(D)** qPCR analysis of *Il6*, *Il1b* and *Tnf* mRNA expression relative to *Gapdh* levels in the DRG at 7 days after SNI or naïve in *Cx3cr1^GFP/GFP^* and *Cx3cr1*^+/+^ mice (n = 5-6). Error bars show the mean ± S.E.M. P values were determined by **(A)** two-tailed Student’s *t* test and **(D)** one-way ANOVA followed by Bonferroni’s post hoc test. *, P < 0.05. Data are representative of at least 2 independent experiments.

## 3. Discussion

Tissue-resident macrophages are abundant in all tissues, whereby they contribute to the maintenance of tissue homeostasis and act as effectors of innate immunity. Among the subtypes of tissue-resident macrophages, sNAMs are distributed along the sciatic nerve and closer to the cell body of primary sensory neurons located in the sensory ganglia where they might be involved in physiological and pathophysiological processes (Wang et al. 2020; Yu et al. 2020; Montague et al. 2018; Kalinski et al. 2020; Old et al. 2014; Ydens et al. 2020). Evidence indicates that after peripheral nerve injury macrophage population increases in the sensory ganglia and accounts for neuropathic pain development and neuroregeneration, through the production of pronociceptive mediators, such as cytokines and chemokines (Yu et al. 2020; Santa-Cecília et al. 2019). The nerve injury-induced sNAMs expansion in the sensory ganglia has been ascribed to the infiltration of blood monocytes. Nevertheless, this hypothesis is still debated and additional evidence to clarify this process needs to be obtained. In this study, by using different experimental approaches, we provided strong evidence that after SNI, a well-known model of peripheral nerve injury, the number of macrophages that increase in the DRG is mainly due to the proliferation of CX3CR1^+^ resident macrophages and does not rely on the infiltration of CCR2^+^ blood monocytes. Additionally, we showed that CX3CR1 signaling on sNAMs mediates their proliferation and activation, stimulating the production of TNF and IL-1b cytokines.

It was long considered that after peripheral nerve injury the increase of inflammatory mediators in the DRG is associated with the infiltration of peripheral blood monocytes, assumed from the increased expression of some markers such as IBA1, CD11b, ED1, CD68, and/or F4/80 (Liu et al. 2010; D. Kim, You, et al. 2011; Huang et al. 2014; Kallenborn-Gerhardt et al. 2014; Luo et al. 2019). In order to dispute this hypothesis, we initially characterized the DRG macrophages based on the expression of CCR2 and CX3CR1 receptors which can be used to distinguish two major populations of macrophages/monocytes (Gordon and Taylor 2005). We took advantage of the transgenic reporter mice *Cx3cr1^GFP/+^/Ccr2^RFP/+^* to trace the fate of tissue-resident macrophages (CX3CR1^+^) and monocytes recruited from the blood to inflamed tissue (CCR2^+^) after peripheral nerve injury. The same strategy has been used to distinguish peripheral monocyte infiltration in the central nervous system and resident microglia in different models of diseases (Mizutani et al. 2012; Yamasaki et al. 2014; Chen et al. 2020). Contrary to previous studies, the number of CCR2^+^ monocytes remained constant in the DRGs after induction of SNI, despite the significant increase in the number of CX3CR1^+^ resident macrophages. These results initially could indicate that blood CCR2^+^ monocytes fail to infiltrate the sensory ganglia after peripheral nerve injury. In fact, our parabiosis experiments support the idea that blood-derived monocytes are not recruited to the DRG after SNI. Furthermore, the parabiosis experiments also revealed that, in fact, no leukocyte subtype is able to significantly infiltrate the sensory ganglia after peripheral nerve injury, similar to how we and others have been demonstrated for the spinal cord (Gu et al. 2016; Guimarães et al. 2019). Even though the peripheral nerve injury did not trigger an infiltration of circulating CCR2^+^ monocytes into the DRG, few of these cells were observed in the tissue parenchyma under homeostasis. These data are in line with recent findings demonstrating that BM-derived monocytes may contribute to a modest subset of PNS macrophages at steady-state. Additionally, the maintenance of DRG-resident macrophages can be slowly replaced by CCR2^+^ monocytes, while the majority of sNAMs arise from embryonic precursors and must be able to proliferate and self-renew (Wang et al. 2020).

One important question that arises from these data is why peripheral blood-circulating cells are not able to infiltrate the DRG after peripheral nerve injury admitting there is a significant production of proinflammatory mediators in the tissue? Although the reasons are not immediately apparent, the possible explanation is the blood dorsal root ganglion barrier, formed by the perineurium and endoneurial blood vessels. This barrier protects and maintains the PNS in an appropriate physicochemical environment (Reinhold and Rittner 2020). The perineurium is a thick layer of connective tissue whose cells have a non-polarized architecture and are interconnected by tight junctions (TJ), gap junctions, and adherens junctions (AJ), similar to the composition of the CNS blood-brain barrier. The endoneurial vessels are composed of a network of arterioles, venules, and non-fenestrated capillaries. The endothelial cells that create this vascular network are also sealed by TJ, but more permeable than the perineurium, since there must be a controlled exchange between blood and nerve to allow neural nutrition (Reinhold and Rittner 2017). While these barriers may limit the infiltration of circulating monocytes into the DRG under homeostasis, it remains unclear how peripheral nerve damage induced by SNI can affect its integrity. Previous studies suggest that after crush injury there may be a loss and recovery of the blood dorsal root ganglion barrier junction, associated with the expression of intercellular junctional proteins (Hirakawa et al. 2003). Although immune cells seem to be unable to infiltrate the cell-body-rich area of the DRGs after peripheral nerve injury, there is recent evidence of accumulation of leukocytes into the dorsal root leptomeninges that recover the sensory ganglia (Du et al. 2018). Nevertheless, additional studies will be necessary to elucidate leukocyte trafficking into these regions after peripheral nerve injury.

Our study further showed that SNI-induced increase of sNAMs in the sensory ganglia is a consequence of the local proliferation of CX3CR1-resident macrophages. Our data showing an increase in the expression of Ki67, a classical marker of cell proliferation, in CX3CR1^+^ cells before the expansion of this population, is an important finding to support this conclusion. Additionally, a recent study that analyzed the single-cell transcriptome of DRG cells after peripheral nerve injury also found increased expression of proliferation genes in the macrophages population (Avraham et al. 2020). Nevertheless, it is still unclear how the peripheral nerve injury leads to the distal proliferation/activation of sNAMs seeded in the DRG. Recent findings indicate that constant stimulation of neurons following peripheral nerve injury results in CX3CL1 production in the spinal cord, which in turn induces activation/proliferation of local microglia(Verge et al. 2004; Peng et al. 2016; Clark et al. 2007). Here, we have shown that CX3CL1/CX3CR1 signaling seems to be also involved in sensory ganglia macrophages activation/proliferation. In fact, in the DRG, the CX3CR1-proliferating macrophages are in close contact with the cell body of sensory neurons, which constitutively express the membrane-bound CX3CL1 (K.W. Kim, Vallon-Eberhard, et al. 2011; Huang et al. 2014). Moreover, after peripheral nerve injury, membrane-bound CX3CL1 is reduced in the cell bodies of sensory neurons, suggesting their release and action in the sNAMs (Zhuang et al. 2007). Since the increase in the number of sNAMs was only partially reduced in CX3CR1 deficient mice, it is plausible that other signaling pathways are involved in the activation/proliferation of sNAMs in the DRGs after peripheral nerve injury. One possibility that has been recently explored is the CSF1-CSF1R signaling. In fact, injured neurons produce and release CSF-1 that in turn promotes sNAMs expansion in sensory ganglia through CSF1R activation (Guan et al. 2016; Yu et al. 2020).

Like classic cells in the innate immune system, sNAMs in the sensory ganglia also express Toll-like receptors (TLRs) and nucleotide-binding cytoplasmic oligomerization (Nod)-like receptors (NLRs). Previous studies indicate that activation of sNAMs in the sensory ganglia, after peripheral nerve injury, depends on downstream signaling generated by the activation of TLR2, TLR4, and TLR9 (Shen et al. 2017; D. Kim, You, et al. 2011; Luo et al. 2019). We also recently demonstrated that NOD2 deficiency prevented the increase in the number of sNAMs in the DRGs after SNI (Santa-Cecília et al. 2019). Considering the involvement of PRRs in the activation/proliferation of DRG-resident macrophages, future studies will be necessary to clarify how these cells recognize or respond to a peripheral nerve injury, which is assumed to be a sterile condition.

Finally, we also address the importance of sNAMs for the production of pro-inflammatory cytokines, especially TNF, IL-1b, and IL-6, which have been described as upregulated in the sensory ganglia after peripheral nerve injury (Ji, Xu, and Gao 2014). Although there is a consensus that peripheral nerve injury triggers the up-regulation of these cytokines in the sensory ganglia, their cellular source is not completely characterized and there are discrepancies in the literature. We provided evidence by using different approaches that allow us to strongly indicate that, at least, TNF and IL-1b, are produced mainly by sNAMs. The production of IL-1b and TNF by sNAMs in the sensory ganglia has been recently implicated in the pathophysiology of neuropathic pain (Santa-Cecília et al. 2019; Yu et al. 2020). Nevertheless, our data indicate that sNAMs are not the source of IL-6 after peripheral nerve injury. This is in agreement with some reports indicating that sensory neurons start to produce IL-6 after different nerve injuries (Hu et al. 2020). Furthermore, our data showing that CX3CR1 signaling is not involved in the IL-6 production after peripheral nerve injury, further support the independence of its production by sNAMs.

In summary, the present study elucidates that the increase of sNAMs in the DRG triggered by peripheral nerve injury is a result of resident macrophages proliferation, and does not depend on the infiltration of blood-derived CCR2^+^ monocytes. Furthermore, CX3CR1 signaling in sNAMs mediates their activation/proliferation, as well as the production of pro-inflammatory mediators. In conclusion, our findings might be useful to explore the modulation of sNAMs proliferation in conditions related to peripheral nerve injuries such as neuropathic pain and nerve regeneration.

## 4. Methods

### 4.1 Animals

For all experiments we use 7-10-week-old males, unless specified in the text. C57BL/6 wild-type (WT) were purchased from Jackson Laboratory, bred and raised in house. *Ccr2^RFP/RFP^* mice (Saederup et al. 2010), and *Cx3cr1^GFP/GFP^* mice (Jung et al. 2008). *Ccr2^RFP/+^*-*Cx3cr1^GFP/+^* mice were generated by crossbreeding *Ccr2^RFP/RFP^* mice with *Cx3cr1^GFP/GFP^* mice. We also used hemizygous transgenic mice expressing eGFP, C57BL/6-(Tg[CAG-EGFP]), under the control of the chicken-actin promoter and cytomegalovirus enhancer (Okabe et al. 1997). Local colonies of transgenic mice were then established and maintained on a C57BL/6 background at the animal care facility of the Ribeirao Preto Medical School, University of Sao Paulo. Food and water were available *ad libitum*. Animal care and handling procedures were under the guidelines of the International Association for the Study of Pain for those animals used in pain research and were approved by the Committee for Ethics in Animal Research of the Ribeirao Preto Medical School— University of São Paulo (Process number 002/2017).

### 4.2 Spared nerve injury model

Spared-nerve injury (SNI) was used as a model of a distal peripheral nerve injury. Briefly, animals were anesthetized with isoflurane, and the sciatic nerve and its 3 terminal branches were exposed. The tibial and common peroneal branches were ligated using 5-0 silk and sectioned distally, whereas the sural nerve remained intact, as previously described (Decosterd and Woolf 2000). Finally, the muscle and skin were sutured in 2 layers.

### 4.3 HSV-1 infection

Mice were anesthetized with isoflurane and then the mid flank and right foot were clipped and depilated with a chemical depilatory (Veet Hair Remover; Reckitt Benckiser). Three days later, HSV-1 (2 x 10^5^ PFUs in 20 ul) was inoculated on the skin of the right hind paw (5×5mm) after scarification with sandpaper. The virus was applied directly to the scarified area (J.R. Silva et al. 2017).

### 4.4 Quantitative real-time RT-PCR

At the indicated time points after nerve injury (SNI) or naive mice were terminally anesthetized with xylazine ketamine and then transcardially perfused with phosphate-buffered solution (PBS 1x). The dorsal root ganglia (L4-L5) ipsilateral to the lesion were collected and rapidly homogenized in 500 *μ*l of TRIzol solution (Thermo Fischer Scientific) reagent at 4^◦^C. Then, total cellular RNA was purified from the tissue according to the manufacturer’s instructions. The purity of total RNA was measured by a spectrophotometer using the wavelength absorption ratio (260/280 nm), that was between 1.8 and 2.0 for all preparations. The obtained RNA samples were reverse-transcribed with High Capacity Kit (Thermo Fischer Scientific). Real-time PCR was performed using specific primers for the mouse genes *Aif1*, *Cx3cr1, Il1b, Tnf, Il6, Cxcl1, Csf1r, and Kcnj1* (Table 1). The levels of each gene were normalized to the expression of *Gapdh*. Reactions were conducted on the Step One Real-Time PCR System using the SYBR-green fluorescence system (Applied Biosystems, Thermo Fisher Scientific, Waltham, MA, USA). The results were analyzed by the quantitative relative expression 2^−ΔΔCt^ method as previously described (Livak and Schmittgen 2001).

### 4.5 Immunofluorescence analyses and quantification

At the indicated times after nerve injury, mice were terminally anesthetized with xylazine ketamine and transcardially perfused with PBS 1x. Following by 4% paraformaldehyde (PFA) in 0.1 M PBS, pH 7.4 (4°C). After the perfusion, the dorsal root ganglia (L4-L6) were dissected, post-fixed in PFA for 2 h, and then bathed in 30% sucrose overnight. The DRGs were covered with tissue Tek (Electron Microscopy Sciences, 62550–01) and sections were cut (15 *μ*m) in a cryostat (Leica Biosystems, CM3050S). Then, the sections were incubated in blocking buffer, 2%BSA and 0.3%Triton X-100 in PBS. After 1 h, the sections were incubated overnight at 4°C with polyclonal primary antibody for ionized calcium-binding adapter molecule 1 (IBA-1; 1:400; Wako Chemicals, Richmond, VA, USA) or antibody for proliferation marker (Ki-67; 1:400; Abcam, Cambridge, UK). The sections were washing with PBS and incubated with the appropriate secondary antibody solution for 2h at room temperature (IgG conjugated Alexa Fluor 594 and/or 647; 1:800; Invitrogen, Carlsbad, CA). The sections were washed with PBS and mounted with coverslips adding Aqueous Mounting Medium, Fluoroshield with DAPI (Sigma-Aldrich). For the evaluation of CX3CR1 and CCR2 expression, the aforementioned genetically modified *Ccr2^RFP/+^*: *Cx3cr1^GFP/+^* mice were used.

The DRG sections were acquired on Zeiss LSM780 confocal microscope. Images were processed in FIJI package for ImageJ software and the quantification of macrophages was performing using the cell counter. For all measurements, 3 sections of each DRG (L4 or L5) were analyzed, and the results were averaged to generate the value for a single mouse.

### 4.6 Flow Cytometry and cell sorting

DRGs (L3-L5) were collected from naive or SNI mice and incubated in a solution of 1 ml of RPMI 1640 medium with 2 mg/ml of collagenase type II (Gibco) for 30 minutes at 37^◦^C. After digestion, the DRGs were mechanically grinded through 40-*μ*m cell strainers and the cell suspension was washed with PBS 1x. The cells obtained were resuspended in PBS 1x containing specific monoclonal antibodies against surface markers for 10 min at room temperature. Dead cells were excluded by Fixable Viability Dye (Catalog number 65-0865-14, Thermo Fisher Scientific, 1:3000). The following monoclonal antibodies were used: anti-CD45-BV421 (Clone 30-F11, BD Biosciences, 1:350), anti-CD11b-FITC (Clone M1/70, BD Biosciences, 1:250), and anti-Ly6G-APC (Clone 1A8, BD Biosciences, 1:250). The sample acquisition was performed by FACSVerse flow cytometry instrument (BD Biosciences, San Jose, CA, USA) and data were analyzed using FlowJo software (Treestar, Ashland, OR, USA).

#### Cell sorting

The DRGs were collected from naive or SNI *Cx3cr1^GFP/+^* mice. As described above, the tissues were digested and filtered through a cell strainer. Samples were then centrifuged and the supernatant was discarded. Cellular pellets were pooled (DRGs from 8 mice), resuspended in a solution containing cell surface markers (CD11b, CD45, and Live/Dead), stained for 10 minutes at room temperature, and then further sorted as macrophages (CX3CR1^+^CD11b^+^) in a FACSAria III sorter. The full-gating strategy used to perform cell sorting from sensory ganglia is depicted in figure S4A. Sorted cells were submitted to RNA extraction, reverse-transcribed with High Capacity Kit (Thermo Fischer Scientific), and analyzed by RT-PCR with a Step One Real-time PCR system as described above (Applied Biosystems).

### 4.7 Parabiosis

Parabiosis was performed as previously described (Kamran et al. 2013). Briefly, 8-week old matched female WT and C57BL/6-(Tg[CAG-EGFP]) mice were co-housed for 2 weeks to reduce stress and then surgically attached for 1 month. Then, they were deeply anesthetized with 1% isoflurane (v/v) and a skin incision was made along the contiguous flanks on the prepared side of each animal. Two animals were paired through the skin, each mouse was sutured to each other, enabling a shared circulation between the two mice. 30 days after the recovery from the parabiosis surgery, mice were subjected to the SNI model and surgically separated 7 days after. Blood exchange was confirmed upon separation by examining GFP^+^ cells in the bloodstream of WT mice by flow cytometry.

### 4.8 Re-analysis of public scRNA-seq data

The scRNA-seq data from mice DRGs was acquired from the Gene Expression Omnibus (GEO) database under the series number GSE139103 (Avraham et al. 2020). The single-cell libraries were generated using GemCode Single-Cell 3′ Gel Bead and Library Kit on the 10X Chromium system (10X Genomics). The dataset contains cells from four animals, of which two naive mice and two with injured DRG. The feature barcode matrix was analyzed using Seurat v3. The cells were filtered according to the criteria: 600-10000 total reads per cell, 500 - 4000 expressed genes per cell, and mitochondrial reads <10%. Clusters were identified using shared nearest neighbor (SNN) based clustering based on the first 30 PCAs and resolution = 0.5. The same principal components were used to generate the t-SNE projections. Differentially expressed genes between samples from naive and injured mice for each cluster were identified using FDR < 0.05 and |avg_log2FC| > 0.25.

### 4.9 Experimental study design (statistics details)

The n sample was determined based on previous publications and/or internal pilot data, to be adequate for statistical analysis and ensured reproducibility. No statistical methods were used to determine the samples size. Additionalty, experimental groups blinded during qualifications. Data are reported as the means ± S.E.M. per group with 4–6 mice. Result analysis was performed by One-way ANOVA followed by the Bonferroni test (for 3 or more groups) comparing all pairs of columns. Alternatively, an unpaired Student’s *t-*test was used to compare 2 different groups. Values of *p <* 0.05 were considered statically significant. Statistical analysis was performed with GraphPad Prism 8 software.

### 4.10 Data availability

All data generated or analyzed during this study are included in the manuscript. Public scRNA-seq data are available in Gene Expression Omnibus (GEO) database under the series number GSE139103 (Avraham et al. 2020).

---

**Appendix 1.**
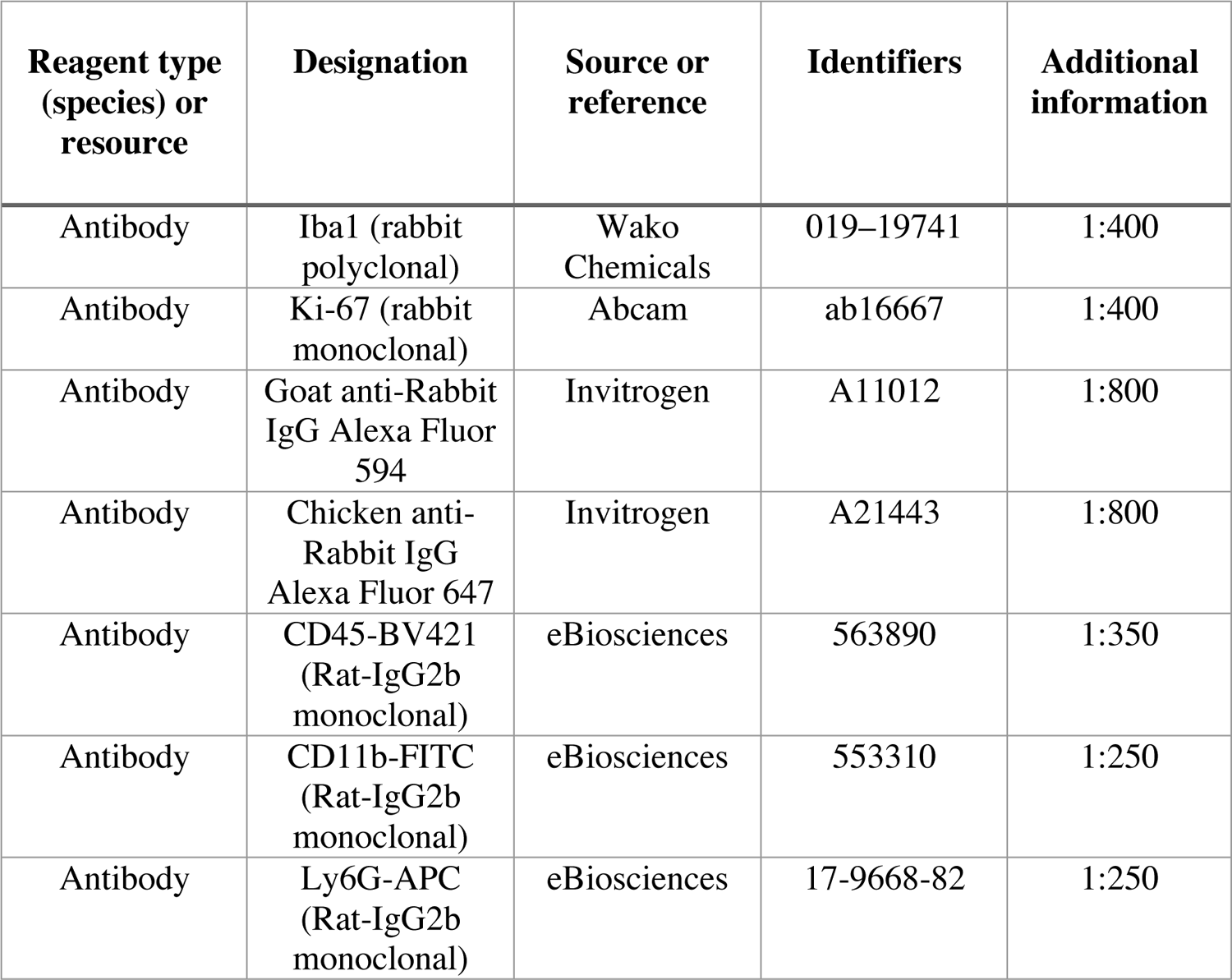
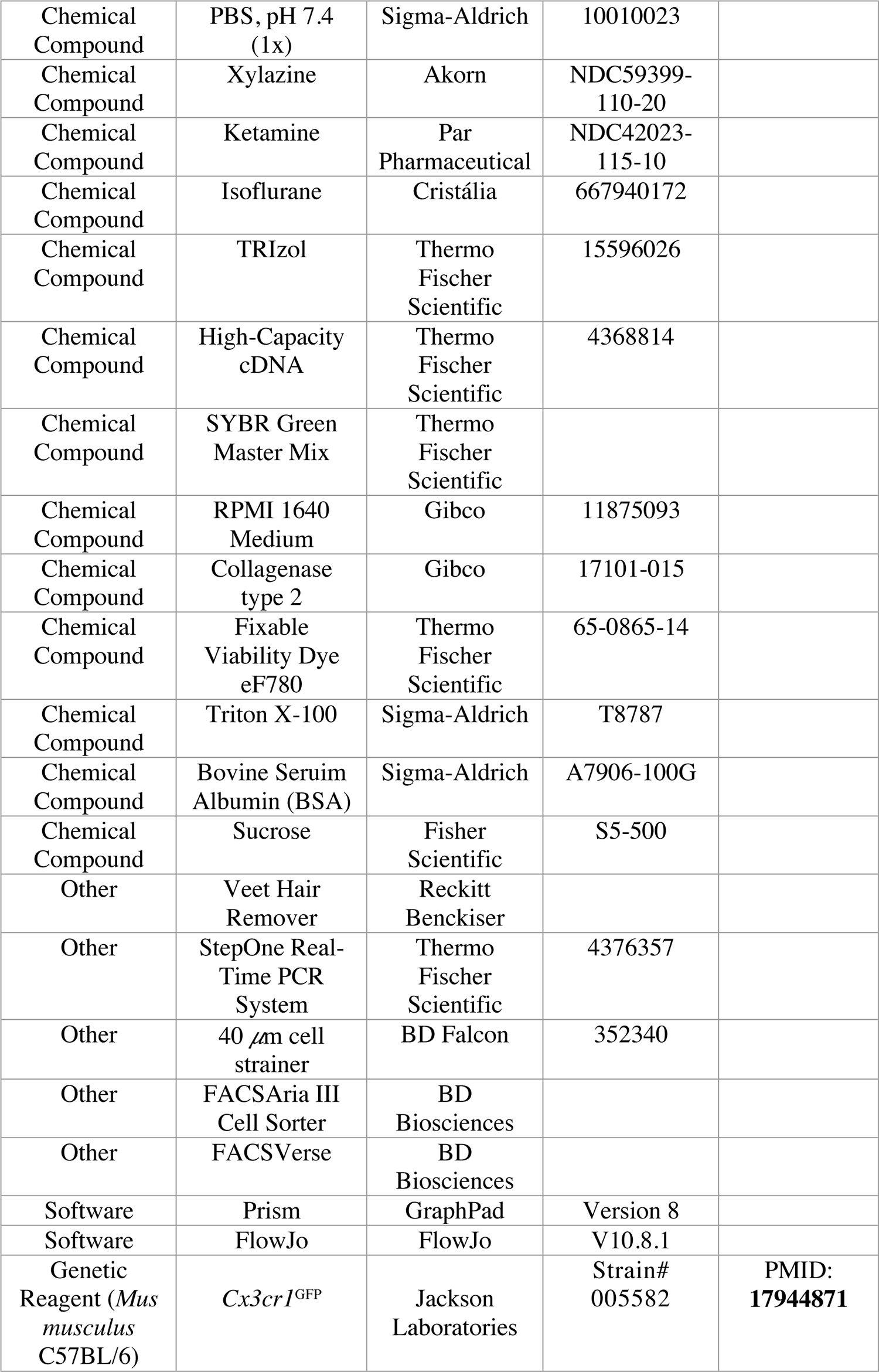
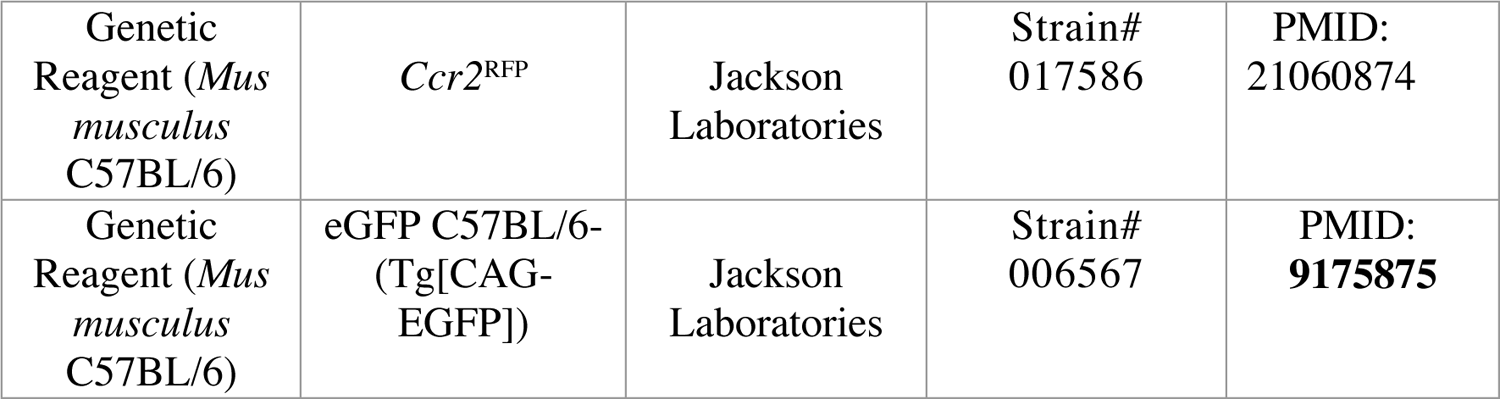
Key resources table

**Appendix 2.**
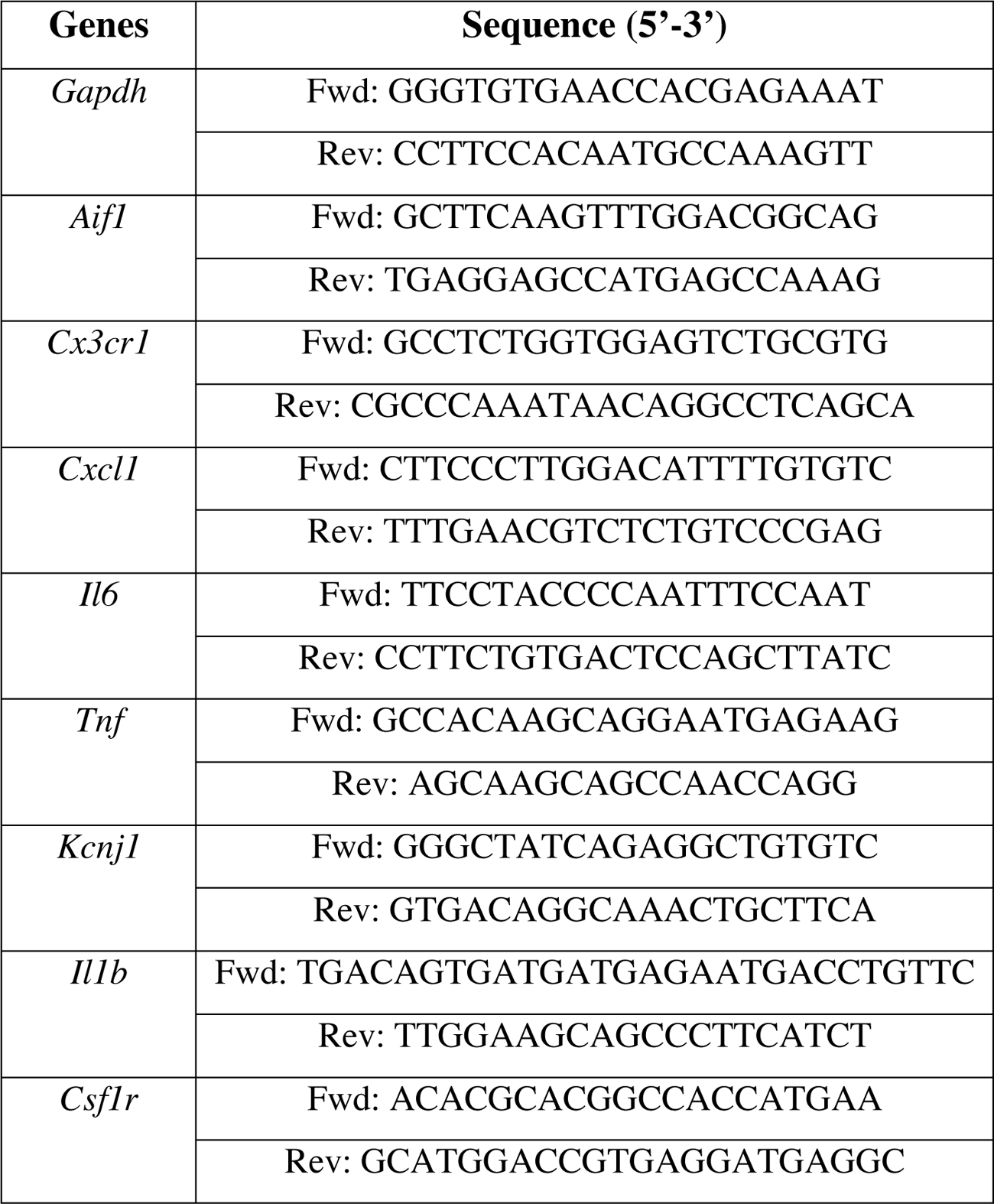
Primers

## ACKNOWLEDGMENTS

The authors gratefully acknowledge the technical assistance of Ieda Santos, Marco Antônio Ribeiro, and Katia Santos for technical assistance. The research leading to these results has received funding from the São Paulo Research Foundation (FAPESP) under grant agreement n◦ 2013/08216-2 (Center for Research in Inflammatory Disease); from Coordenação de Aperfeiçoamento de Pessoal de Nível Superior (CAPES).

---

**Supplementary Figure 1 S1.**
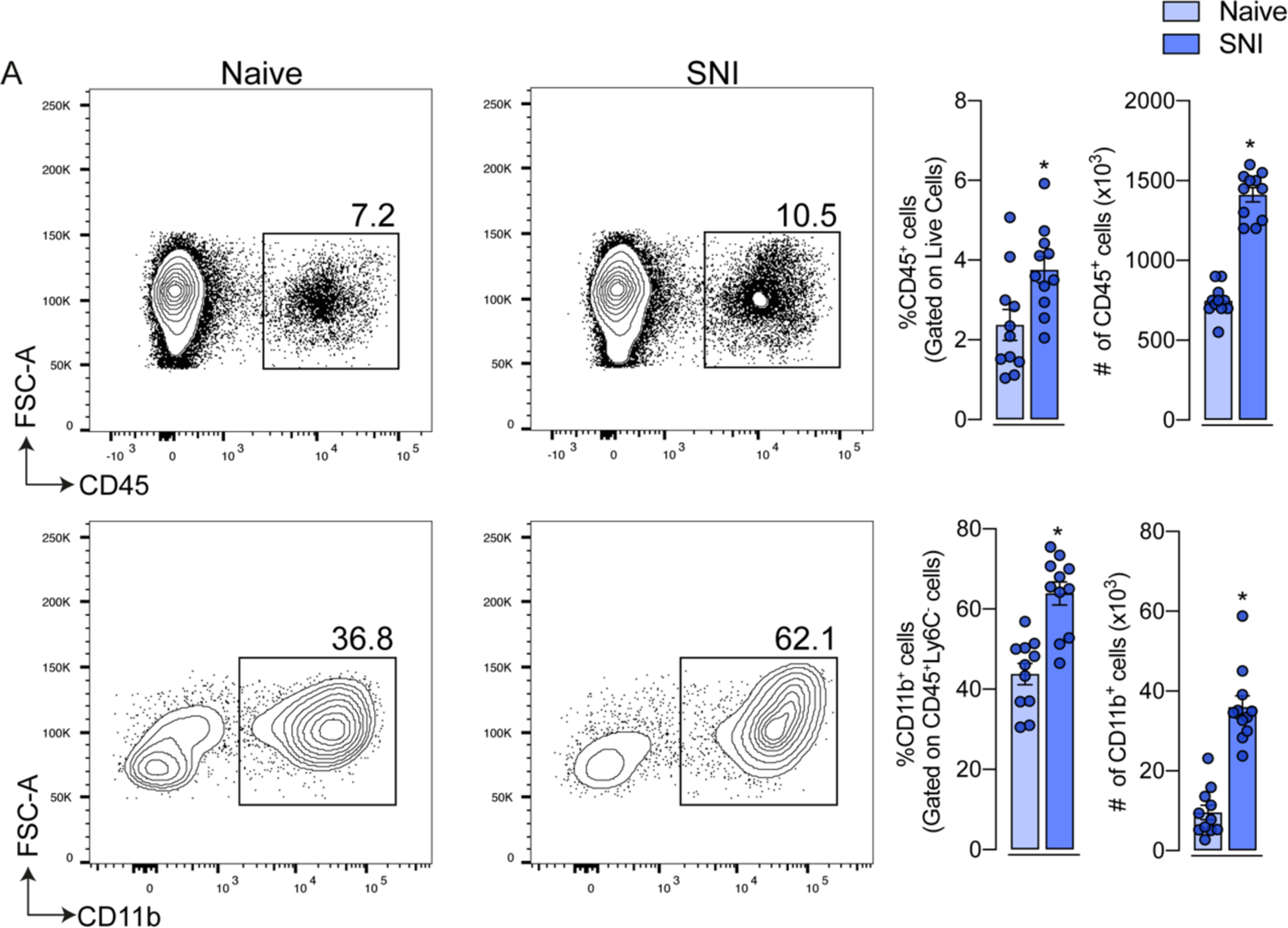
FACS analysis of macrophages population in the DRGs after SNI. (A) Representative dot plots, frequency and absolute number of CD45^+^ and CD11b^+^ Ly6G^-^ cells were evaluated in the DRGs (L3-L5) 7 days after SNI by flow cytometry (n = 10-11). Error bars show mean ± SEM. P values were determined by two-tailed Student’s *t* test. *, P < 0.05; ns, not significant. Data are representative of at least 3 independent experiments.

**Supplementary Figure 2 S2.**
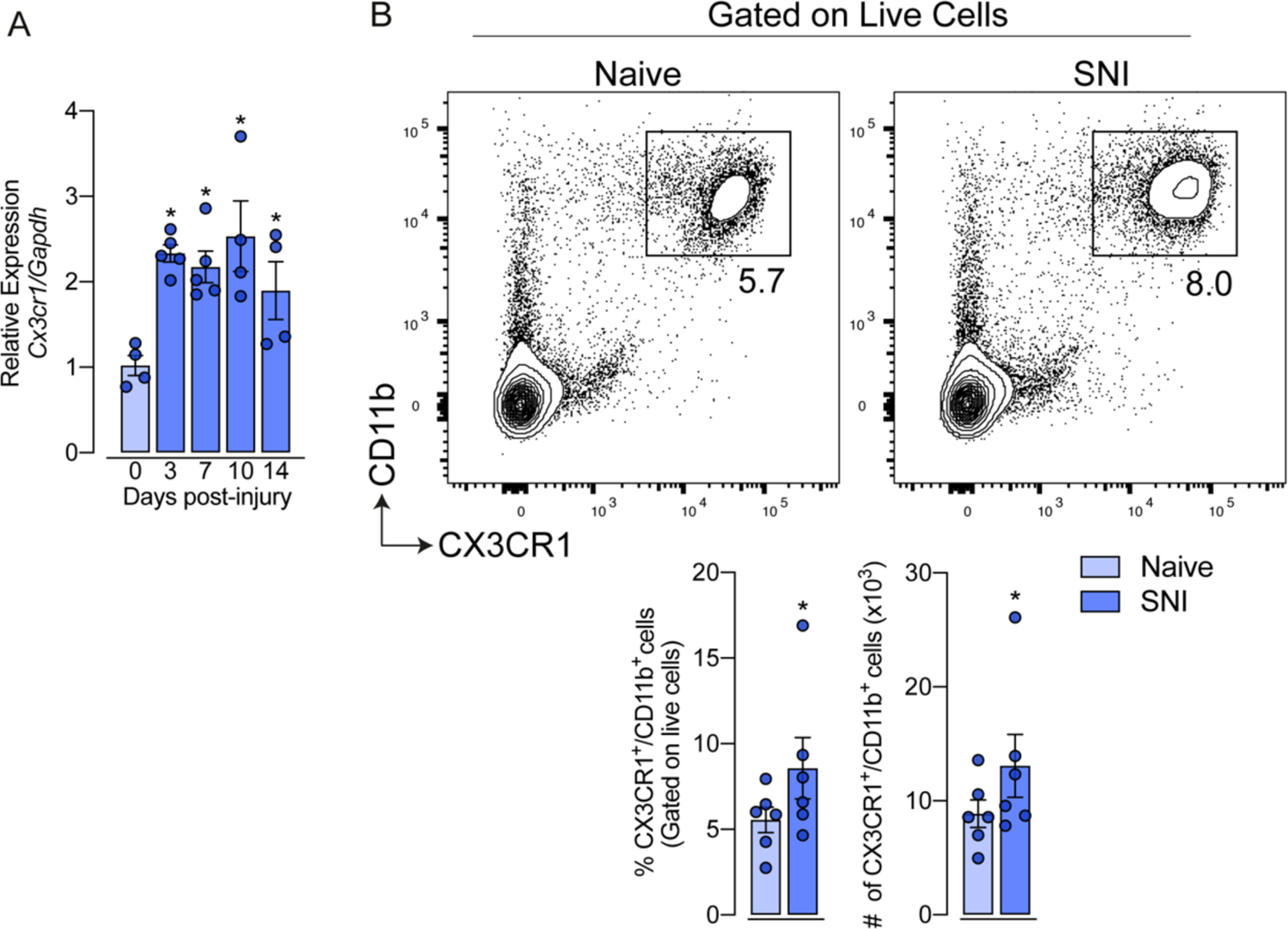
CX3CR1^+^ macrophages number increase in the DRGs after SNI. (**A**) qPCR analysis of *Cx3cr1* mRNA expression relative to *Gapdh* levels in the DRG at 7 days after SNI or naive in C57BL/6 mice (n = 4-5). **(B)** Representative dot plots, frequency and absolute number of CX3CR1^+^CD11b^+^ cells were evaluated in the DRG from C57BL/6 mice at 7 days after SNI by flow cytometry (n = 6). Error bars show the mean ± S.E.M. P values were determined by **(A)** one-way ANOVA followed by Bonferroni’s post hoc test and **(B)** two-tailed Student’s *t* test. *, P < 0.05. Data are representative of at least 3 independent experiments.

**Supplementary Figure 3 S3.**
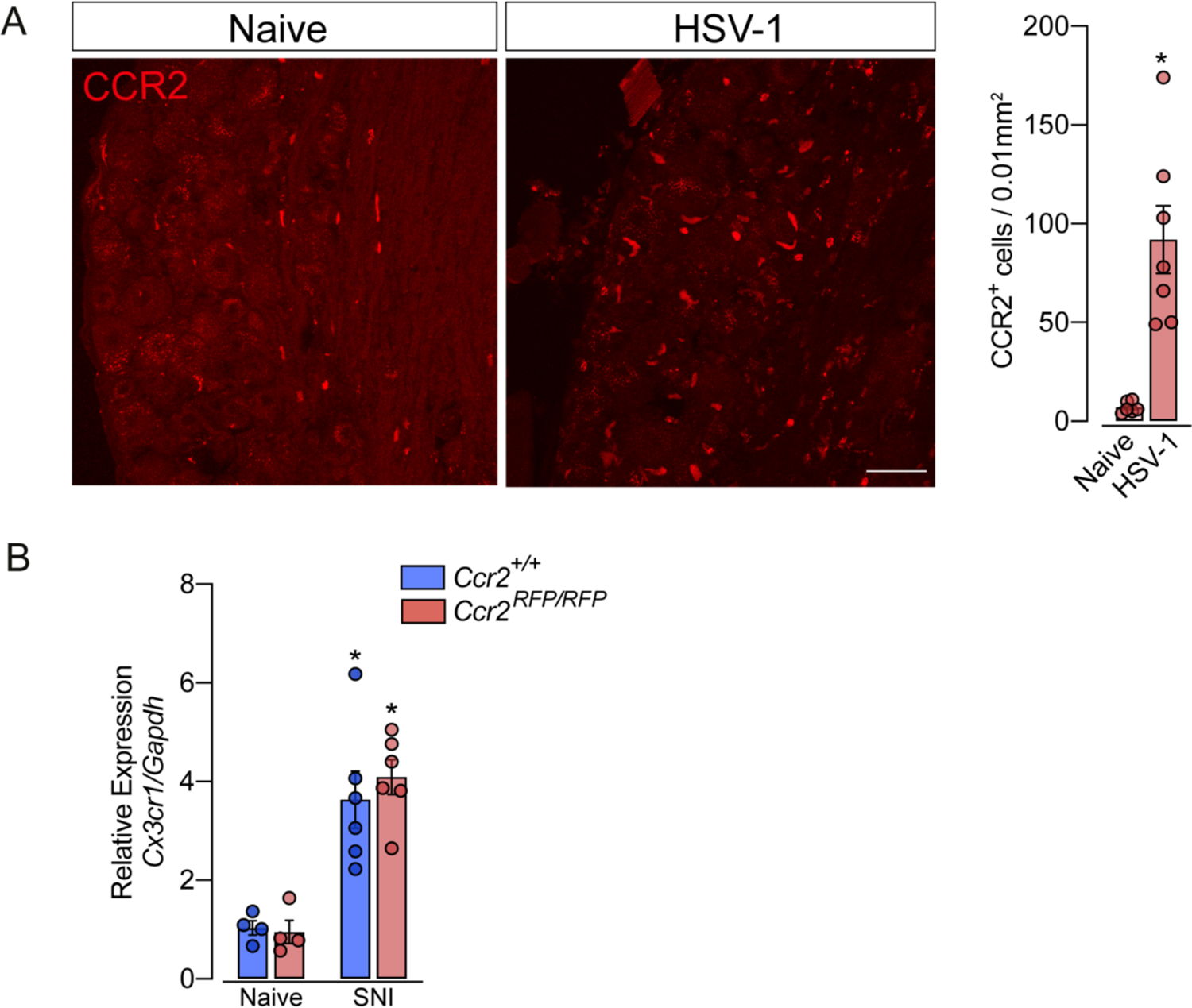
CCR2^+^ monocytes/macrophages in the DRGs after HSV-1 peripheral infection. (A) Representative confocal images of DRGs from *Ccr2^RFP/+^* mice at 7 days after HSV-1 induction. CCR2+ cells are shown in red (n = 7). Scale bars: 50 μm. **(B)** qPCR analysis of *CX3CR1* mRNA expression relative to *Gapdh* levels in the DRG at 7 days after SNI or naïve in *Ccr2^RFP/RFP^* and *Ccr2*^+/+^ mice (n = 4-6). Error bars show the mean ± S.E.M. P values were determined by **(A)** two-tailed Student’s *t* test and **(D)** one-way ANOVA followed by Bonferroni’s post hoc test. *, P < 0.05. Data are representative of at least 2 independent experiments.

**Supplementary Figure 4 S4.**
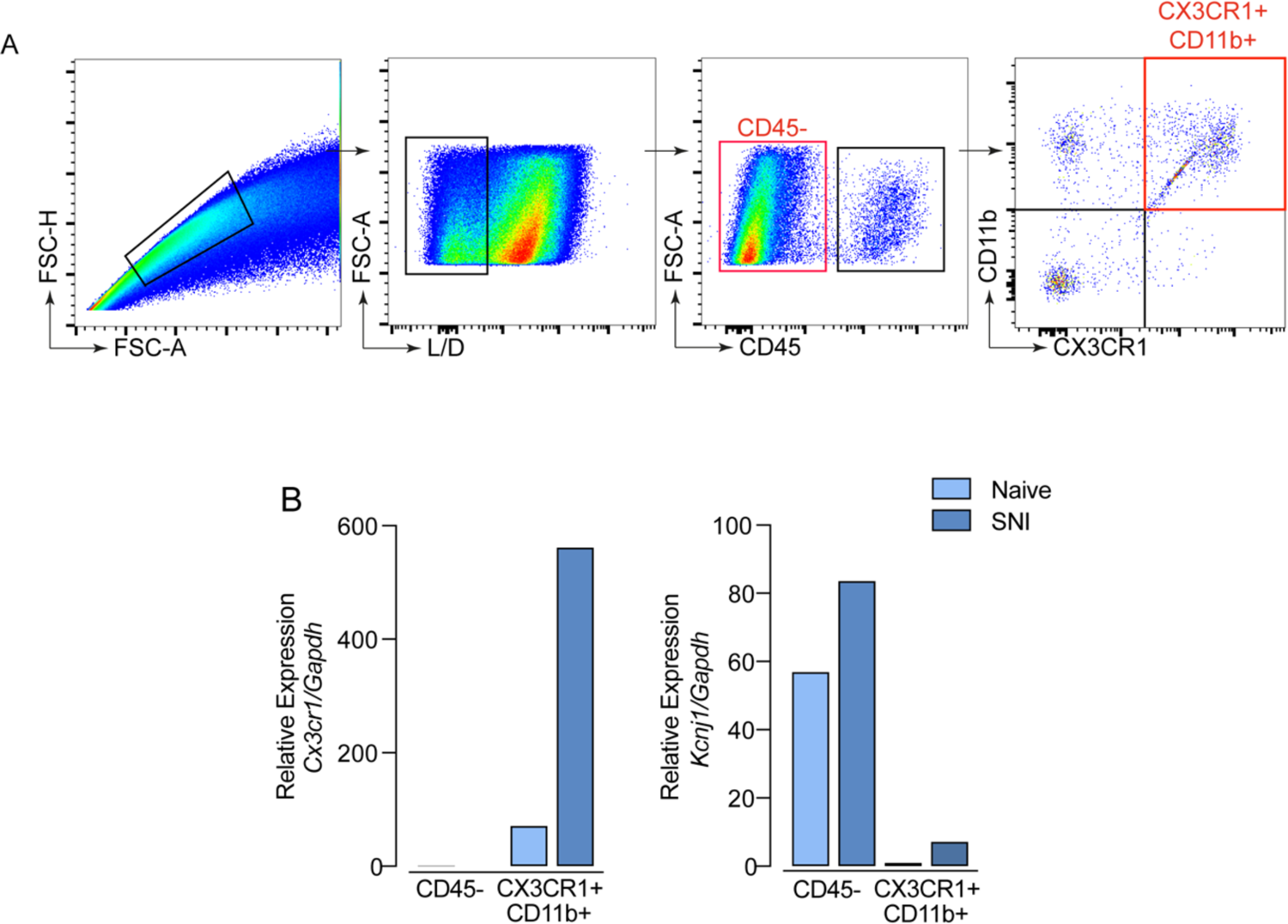
Representative gating strategies for flow cytometry analysis and cell sorting. (A) Cells suspension isolated from DRGs of *Cx3cr1^GFP/+^* mice (Naive and 7 days after SNI; n= 6 pooled) were used for cell sorting strategy. FSC-H/FSC-A preliminary gate was performed for all cytometry analysis to exclude cell CD45**^+^** debris and cell doublets. Second gate shows gating on viable cells. Next, cells were gated on CD45**^+^** populations. Finally, *Cx3cr1^GFP/+^* within CD11b**^+^** were sorted as well as CD45**^-^** cells. **(B)** Expression of *Cx3cr1* and *Kcnj1* mRNA in *Cx3cr1^GFP/+^* and/or CD45**^-^** sorted cells. Data are representative of at least 3 independent experiments.

## Notes

### Competing Interest Statement

The authors have declared no competing interest.

